# Malaria in pregnancy regulates P-glycoprotein (P-gp/*Abcb1a*) and ABCA1 efflux transporters in the mouse visceral yolk sac

**DOI:** 10.1101/2020.06.27.175018

**Authors:** Lilian M. Martinelli, Klaus N. Fontes, Mila W. Reginatto, Cherley B. V. Andrade, Victoria R. S. Monteiro, Hanailly R. Gomes, Joao L. Silva-Filho, Ana A. S. Pinheiro, Anamaria R. Vago, Fernanda R. C. L. Almeida, Flavia F. Bloise, Stephen G. Matthews, Tania M. Ortiga-Carvalho, Enrrico Bloise

## Abstract

Malaria in pregnancy (MiP) induces intrauterine growth restriction (IUGR) and preterm labor (PTL). However, its effects on yolk sac morphology and function are largely unexplored. We hypothesized that MiP modifies yolk sac morphology and efflux transport potential by modulating ABC efflux transporters. C57BL/6 mice injected with *Plasmodium berghei ANKA* (5×10^5^ infected-erythrocytes) at gestational day (GD) 13.5, were subjected to yolk sac membrane harvesting at GD18.5 for histology, qPCR and immunohistochemistry. MiP did not alter the volumetric proportion of the yolk sac’s histological components. However, it increased levels of *Abcb1a* mRNA (encoding P-glycoprotein) and macrophage migration inhibitory factor (*Mif*-chemokine), whilst decreasing *Abcg1* (P<0.05); without altering *Abca1, Abcb1b, Abcg2, Snat1, Snat2*, interleukin (*Il*)-*1 β* and C-C Motif Chemokine Ligand 2 (*Ccl2*). Transcripts of *Il-6*, chemokine (C-X-C motif) ligand 1 (*Cxcl1*), *Glut1* and *Snat4* were not detectible. ABCA1, ABCG1, breast cancer resistance protein (BCRP) and P-gp, were primarily immunolocalized to the cell membranes and cytoplasm of endodermic epithelium but also in the mesothelium and in the endothelium of mesodermic blood vessels. Intensity of P-gp labeling was stronger in both endodermic epithelium and mesothelium, whereas ABCA1 labeling increased in the endothelium of the mesodermic blood vessels. The presence of ABC transporters in the yolk sac wall suggest that this fetal membrane acts as an important protective gestational barrier. Changes in ABCA1 and P-gp in MiP may alter the biodistribution of toxic substances, xenobiotics, nutrients and immunological factors within the fetal compartment and participate in the pathogenesis of malaria induced-IUGR and PTL.

## Introduction

The harmful effects of malaria in pregnancy (MiP) include high rates of maternal anemia and death, as well as spontaneous abortion, fetal intrauterine growth restriction (IUGR), preterm labor (PTL), low birth weight, fetal/neonatal demise and impaired postnatal cognitive and neurosensory development [1–3]. MiP is characterized by adherence of Plasmodium falciparum-infected erythrocytes to specific syncytiotrophoblast glycosaminoglycans; a condition referred to as “placental malaria” [3,4]. In response, the placental barrier may undergo various adaptive changes that comprise activation and migration of immune cells, alteration of cytokine and chemokine output [2,5], intervillositis, decreased enrichment of syncytiotrophoblast-microvilli and lower expression of placental glucose (GLUT1), amino acid (*SNAT1, SNAT2, Cat1, Lat1* and *4F2hc*), solute carrier (SLC) drug uptake (*Oatp2b1, Oct3, Ent1* and *Ent2*) and nutrient/drug efflux ABC transport systems (ABCA1, BCRP and P-gp) [6–11].

The ATP-Binding Cassette (ABC) transport system is highly expressed in different trophoblastic lineages [12–14]. Its disruption has been associated with IUGR, PTL, pre-eclampsia and chorioamnionitis [13, 15–17]. In this context, reports from different groups have demonstrated that it participates in the adaptive trophoblastic responses to maternal infection [9, 12, 14, 16–18]. Comprised of 50 transporters divided into seven sub-classes ranging from ABCA through ABCG [19], these transmembrane transporters are expressed in different cell types, including those from biological barriers [12]. They are responsible for the efflux of diverse endogenous and exogenous substrates, from one side to the other of the plasm membrane [20]. There are a multitude of endogenous substrates including lipids, phospholipids, cytotoxic oxysterols, amino acids, steroid hormones, folate, metabolites and pro-inflammatory cytokines and chemokines. Exogenous substrates comprise environmental toxins (bisphenol A, ivermectin, pesticides) and clinically relevant drugs (antibiotics, antiretrovirals, antidepressants and synthetic glucocorticoids). ABC transporters are major components of the immune response and mediate nutrient transfer and fetal protection against drugs and environmental toxins that may be circulating in the maternal blood [12,20,21].

The lipid transporters, ABCA1 and ABCG1 (encoded by the *ABCA1* and *ABCG1* genes, respectively), and the multidrug resistance transporters, P-glycoprotein (P-gp/*ABCB1*) and breast cancer-related protein (*BCRP/ABCG2*) are amongst the best-described ABC transporters in the syncitial barrier. ABCA1, P-gp and BCRP are predominantly expressed in the apical membrane of the syncytiotrophoblast, and efflux their substrates from the fetal side into the maternal circulation, whereas ABCG1 is localized in the syncytiotrophoblast’s basolateral membrane and efflux cholesterol into the fetal circulation [12].

*Plasmodium berghei ANKA* (PBA) infection is largely used to mimic malaria infection in mice. In pregnant BALB/c mice, PBA decreases placental *Abca1, Abcb1a* and *Abcb1b* (both encoding P-gp in rodents) and*Abcg2* [11]. Similarly, in pregnant C57BL/6 dams, PBA leads to IUGR, PTL and impairs the placental expression of ABCA1/*Abca1*, P-gp/*Abcb1b* and BCRP/*Abcg2* [9], suggesting that MiP has the potential to increase fetal exposure to a range of clinically relevant substrates capable of profoundly impacting fetal outcome, via dysfunction of ABC transporters in the placental barrier.

The human yolk sac is the major hematopoietic site [22] and is the source of primordial germ cells in early pregnancy [23]. It is estimated that human yolk sac is viable until the 49^th^ day of gestation [24]. Emerging evidence suggests the yolk sac is an important nutrient exchange site between the coelomic fluid and the fetal capillaries present in its mid-mesodermal layer; or alternatively, it delivers nutrients to the embryo primitive gut via the vitelline duct [24–26]. In mice, the visceral portion of the yolk sac involves the embryo amniotic membrane and is functional throughout pregnancy, mediating the transport of critical substances from the mother to the fetus; acting as a syncytiotrophoblast equivalent throughout intrauterine development [27,28]. It is known that human [24] and mouse [24,29] yolk sac express different ABC transport-carriers, suggesting that this membrane acts together with the placenta and other fetal membranes to form an efficient protective barrier during gestation.

Despite the importance of the yolk sac for embryo and fetal development, no studies have investigated the effects of MiP on yolk sac morphology and efflux transport expression. In the present study, we hypothesized that MiP, in a murine model of malaria induced-IUGR and PTL, modifies the yolk sac morphology and efflux transport potential, via modulation of key ABC efflux transporters. Improved knowledge as to how the yolk sac responds to MiP may improve the understanding of the mechanisms by which malaria impacts pregnancy outcome.

## Methods

### Animals

An animal model of severe experimental malaria, which recapitulates many of the features of the human malarial disease in pregnancy, including placental malarial, IUGR, low birth weight and PTL was used as previously described [9]. This model resulted in a 20% rate of PTL in infected dams, however, all analyses were undertaken in the yolk sac of fetuses born at GD18.5 (i.e. term). Briefly, C57BL/6 female mice (8 to 10 week-old) were housed at room temperature (22°C), with a light cycle of 12h light/12h dark and were mated. Animals were maintained on a commercial Nuvilab^®^ CR1 (Nuvilab, PR, Brazil) chow and water ad libitum. Mating was confirmed by the presence of a vaginal plug, and was defined as gestational day (GD) 0.5. Pregnant dams were injected intraperitoneally (i.p.) with 5×10^5^ erythrocytes infected with *Plasmodium berghei ANKA* (PBA group) or with saline (control group) on GD13.5. Euthanasia was performed at GD18.5, using a pentobarbitol overdose (300 mg/kg i.p.), followed by maternal and fetal decapitation. Six PBA-infected dams from our previous cohort [9], which exhibited peripheral parasitemia (approximately 16% of infected erythrocytes) at GD18.5, were included in this study. Approval was obtained from the Institutional ethics committee (CEUA-190/13), registered within the Brazilian National Council for Animal Experimentation Control (protocol number 01200.001568 / 2013-8). All procedures followed the “Principles of Laboratory Animal Care” formulated by the National Society for Medical Research and the U.S. National Academy of Sciences Guide for the Care and Use of Laboratory Animals.

### Volumetric proportion assessment of mouse visceral yolk sac

The visceral yolk sac of PBA-infected and control animals (n=6/group) were dissected and immediately immersed in buffered 4% paraformaldehyde phosphate solution (24h). After fixation, samples were dehydrated in an ethanol series, diaphanized in xylene, and embedded in paraffin wax (Histopar, SP, Brazil). Sections (4μm) were stained with hematoxylin and eosin (H&E). Image acquisition and analysis were performed on a Zeiss Axiolab 1 photomicroscope, coupled with a CCD camera and computer running Zeiss Axiovision (Carl Zeiss, NY, EUA). The histomorphological analysis was undertaken using the Fiji ImageJ 1.0 program (ImageJ, WI, USA). Estimation of relative volume of the different components of the yolk sac (endodermic epithelium, mesodermal connective tissue, mesodermal blood vessels and mesothelium) was undertaken by superimposing yolk sac histological photomicrographs with a grid of equidistant points (measuring 25μm distance between two points). 1000 points coinciding with each of the histological components evaluated were recorded, yielding a variable number of histological sections evaluated per dam; which corresponded to a total average area of 455 mm^2^ per dam. The volumetric proportion (VP) of each histological component was calculated as VP = NP × 100/1000, where NP = number of equivalent points on each histological component [30].

### qPCR

Samples of the visceral yolk sac (n=6/group) were immersed in RNA Later solution (Invitrogen, MA, USA) and frozen at −20°C until further processing. Extraction of total RNA was undertaken using Trizol (Invitrogen) with homogenization conducted using a Tissuelyser LT (Qiagen, Hilden, Germany). Concentration and purity of samples were determined by spectrophotometric analysis (Implen nanofotometer, Munich, Germany). Total RNA (1μg) was reverse-transcribed into cDNA, using the High Capacity cDNA Reverse Transcription kit (Applied Byosistems, CA, USA), according to the manufacturer's instructions. cDNA (4ul) was added to the primer-containing mix (intron-exon spanning) of each gene of interest (Table 1), as well as with the fluorescent DNA interlayer Eva Green (Biotium, CA, USA), in a total volume of 6μL. The quantification of each gene of interest relative to the reference genes, *Ywhaz, Ppib* and *Gapdh* (table 1), was performed in the QuantStudio Real-Time PCR (Applied Biosystems, CA, USA), with the following amplification cycles: initial denaturation at 50oC for 2 min and then at 95°C for 10 min, followed by 40 cycles of denaturation at 95°C for 15 sec. Annealing was performed at 60°C for 30 sec and extension at 72°C for 45 sec. Differences in mRNA gene expression were calculated according to the 2-^ΔΔCT^ method [31] and the assay was considered acceptable when its efficiency ranged from 95% to 105%.

**Table 1.**
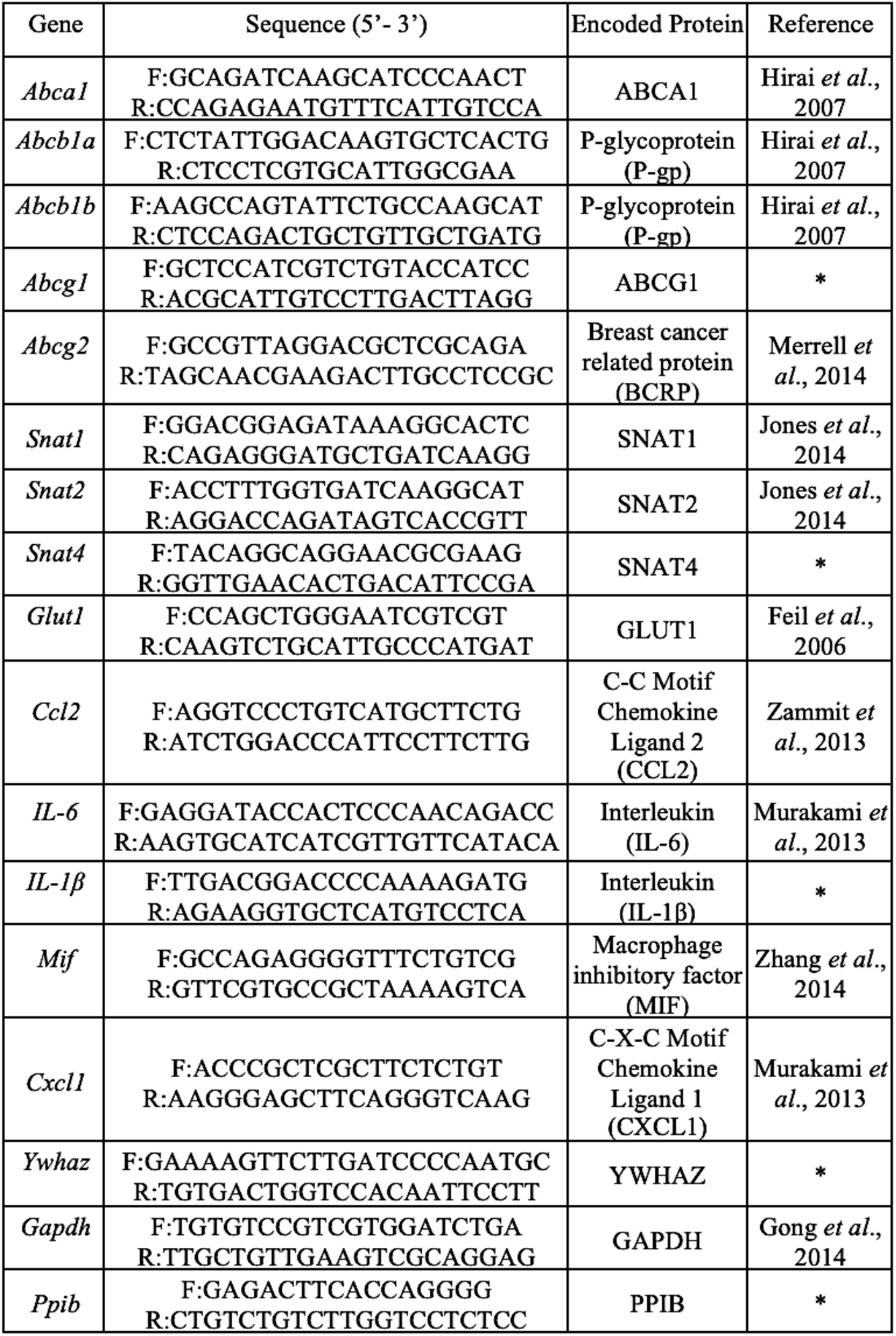
Primers used for qPCR. |

### Immunoh istoch emistry

Yolk sac sections (7μm, n=6/group) were dewaxed and rehydrated. Term mouse placental sections were processed simultaneously, as positive controls. Endogenous peroxidase activity was blocked using the Hydrogen Peroxide Block kit (Springer, Berlin, Germany). Antigen retrieval was performed by heating the sections in Tris-EDTA buffer pH 9, followed by microwave heating in citrate buffer (0.1M, pH 6) allowing to cool on ice (10 min). Blocking was performed by incubation with skimmed milk 10% in PBS (30 min), followed by Protein Block kit (Springer; 30 min). Sections were then incubated overnight with anti-P-gp (mouse monoclonal; Santa Cruz Biotechnology, EUA; 1:500), anti-ABCA1 (mouse monoclonal; Abcam, EUA; 1:100), anti-BCRP (mouse monoclonal; Merck Millipore, GER; 1:200) and anti-ABCG1 (rabbit polyclonal; ThermoFisher Scientific, EUA; 1:100) antibodies. Visualization of protein was performed using the Springer kit following the manufacturer’s instructions. Sections were stained with hematoxylin. Omission of the primary antibody provided negative controls.

Evaluation of the area and intensity of the immunolabeled yolk sac components (endodermal epithelium, connective tissue, endothelium and mesothelium) was performed using a semi-quantitative scoring described previously, with modifications [16,32]. For the immunolabeled area, the scores were: 0) undetectable; 1) 1-25%; 2) 26-50%; 3) 51-75%; and 4) 76-100%. For intensity immunolabeling, graded scores were: 0) no detectable staining; 1) weak; 2) moderate; 3) strong; and 4) very strong intensity. At least 5 fields were evaluated (20x magnification) for each dam. Two operators blinded to the experimental groups performed independent evaluation. An average of the scores for both evaluations was calculated.

### Statistical Method

Values are expressed as mean ± standard error of the mean (SEM) and were analyzed using Prisma program (GraphPad Software Inc., CA, USA). Statistical assessment of volumetric dimensions, qPCR and immunohistochemistry data were undertaken in the yolk sac of fetuses exhibiting the closest placental weight to the mean weight of all placentae from each litter. Thus “n” represents the number of litters [9,33]. After confirming the non-parametric distribution of the data and exclusion of outliers (Grubbs’ test), differences between PBA-infected and control groups were assessed by non-parametric Mann-Whitney test, when comparing two variables; and by Kruskal Wallis test, followed by Dunn’s post-test, when comparing more than two variables. Statistical significance was considered when p <0.05.

## Results

### Malaria does not affect visceral yolk sac histomorphological parameters

The wall of the visceral yolk sac in both PBA-infected and control animals exhibited its three typical layers: an outer (uterine facing) endodermic epithelium, a blood vessel-enriched mesodermal mid-layer and an inner (amnion facing) mesothelial layer. There were no visible differences in gross morphology of these layers between infected and control groups (Figure 1A and B). Additionally, MiP did not affect the volumes of the histological components of the yolk sac, comparing PBA-infected and control groups (Figure 1A). Of note, constitutive endodermic epithelium comprised approximately 66% of the total yolk sac cell number, whereas the constitutive mesothelium formed approximately 5% of the total number of yolk sac cells (Figure 1C).

**Fig. 1.**
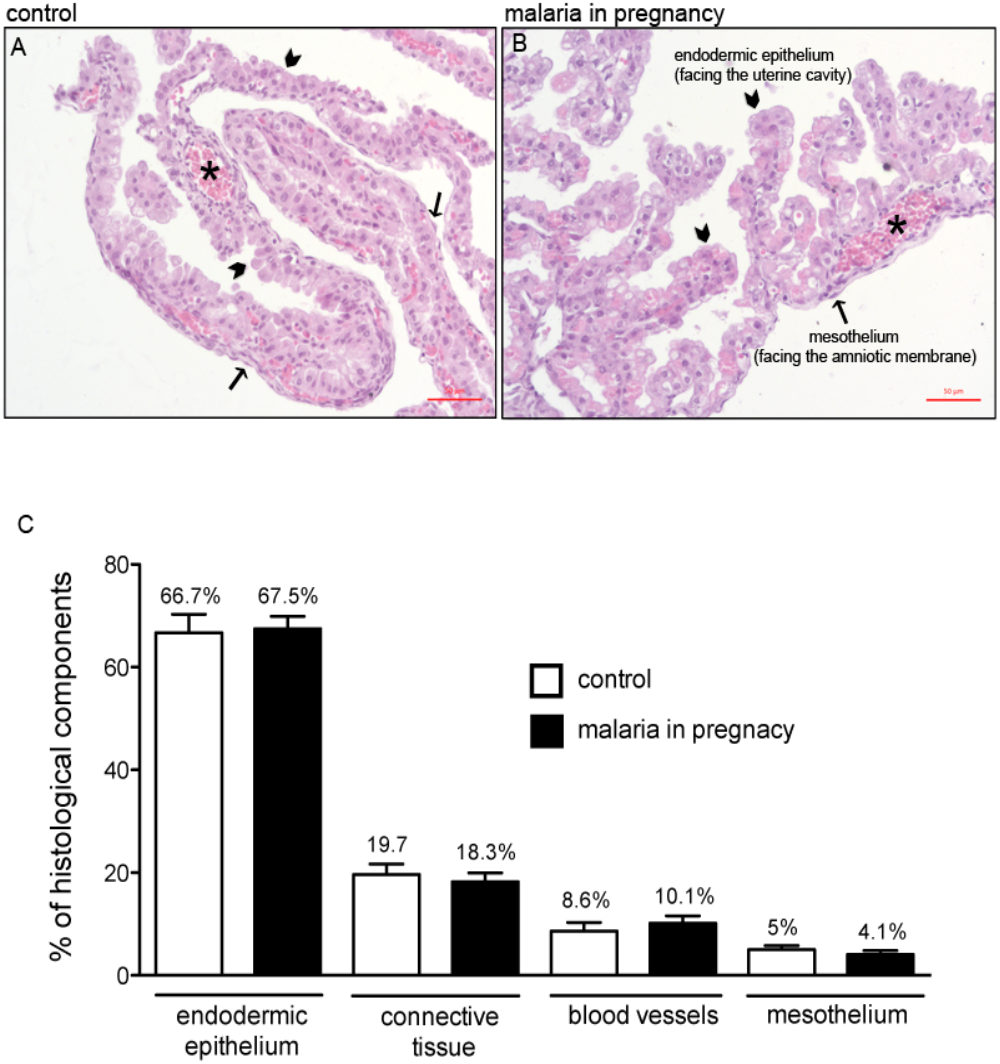
Malaria in pregnancy (MiP) does not affect visceral yolk sac morphology and morphometry. Yolk sac photomicrographs (HE) of **(A)** control and *Plasmodium berghei ANKA* (PBA) infected dams. **C)** Volumetric proportion of the yolk sac histological components from control and PBA-infected dams at GD18.5. Arrowheads = endodermic epithelium; thin arrows = mesothelium; * = mesodermic blood vessels. Statistical differences were tested by Kruskal Wallis test, followed by Dun’s post-test. Data are presented as mean ± SEM (n=6/group). Magnification bars represent 50 μm. 1 % of structures found were classified as artifacts.

### Malaria modifies expression of ABC efflux transporters and pro-inflammatory factors in the yolk sac

mRNA of all ABC transporters evaluated (*Abca1, Abcb1a, Abcb1b, Abcg1* and *Abcg2*) was detected in the mouse visceral yolk sac at term (GD18.5) in both control and in PBA-infected experimental groups. *Abcb1a* mRNA was up-regulated, whereas *Abcg1* was down-regulated in the yolk sac of infected animals, compared to controls (p<0.05, Figure 2). No changes in *Abca1, Abcb1b* and *Abcg2* were observed. Analysis of other active transmembrane (intake) transporters in the yolk sac revealed that *Snat1* and *Snat2* mRNA levels were unaffected by MiP; *Snat4* and *Glut1* transcripts were undetectable in both experimental groups.

**Fig. 2.**
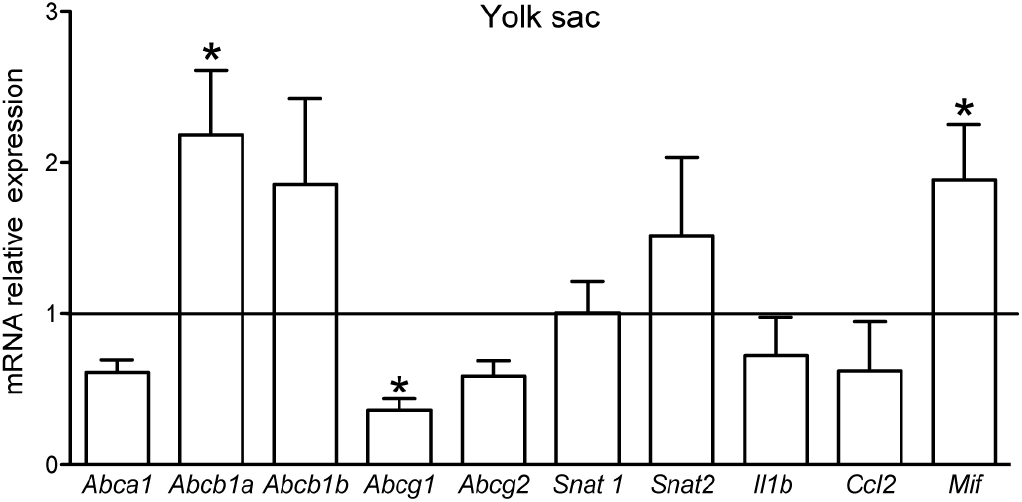
Malaria in pregnancy (MiP) modifies the yolk sac gene expression of specific ABC efflux transporters and pro-inflammatory factors. Relative mRNA expression of selected ABC (*Abca1, Abcb1a, Abcb1b, Abcg1* and *Abcg2*) and nutrient (*Snat1* and *Snat2*) transporters, as well as selected cytokines and chemokines (*Il-1β, Ccl2* and *Mif*) in the yolk sac from *Plasmodium berghei ANKA*-infected dams at GD18.5. Transcripts *of Il-6, Cxcl1, Glut1* and *Snat4* were under detectible limits. Statistical differences were tested by Mann Whitney test. *p<0.05. Data are presented as mean ± SEM (n=6/group).

At the level of the pro-inflammatory genes, we detected increased levels of the macrophage migration inhibitory factor (the *Mif*-chemokine) mRNA, while interleukin (*Il*)-*1 β* and C-C Motif Chemokine Ligand 2 (*Ccl2*) mRNA remained unchanged. Transcripts of *Il-6* and chemokine (C-X-C motif) ligand 1 (*Cxcl1*, a human Il-8 analog) were not detectable in the yolk sac (Figure 2).

### Malaria in pregnancy modifies P-gp and Abca1 protein in the yolk sac

We next evaluated the effects of PBA infection on protein localization and expression (semi-quantitative) of the ABC transporter that exhibited altered gene expression following infection. P-gp was enriched in the apical membrane and cytoplasm of the outer endodermic epithelium and inner mesothelial cells. The endothelium of blood vessels, localized in the mesodermal mid-layer, exhibited faint or no P-gp labeling, whereas the mesodermal layer connective tissue was completely negative for P-gp (Figure 3A and B). In PBA-infected pregnancies, distribution of yolk sac P-gp was similar to controls, however, semi-quantitative analysis revealed increased P-gp protein in the plasma membranes of the endodermic epithelium and mesothelium (p<0.05), with no changes in the total area of Pgp immunolabeling (Figure 3F and G).

**Fig. 3.**
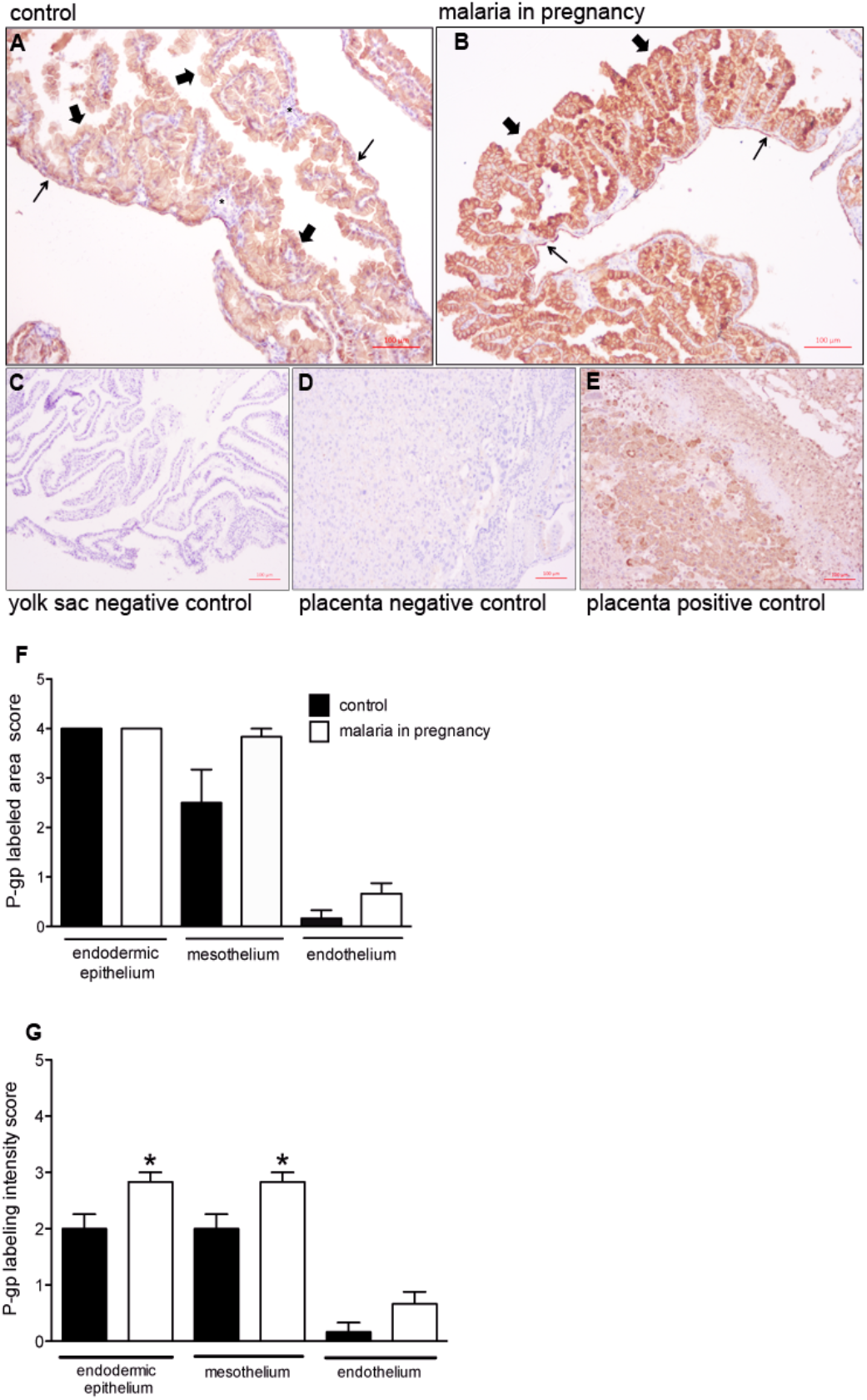
P-glycoprotein (P-gp) is localized in distinct cellular barriers of the murine yolk sac and is upregulated by malaria in pregnancy (MiP). Representative immunohistochemistry images from murine yolk sac sections of control **(A)** and **(B)** *Plasmodium berghei ANKA-infected* dams at GD18.5. Yolk sac **(C)** and placental (**D)** negative controls. **(E)** placental positive control showing P-gp immunoreactivity in labyrinthine and junctional zone cells. Semiquantitative evaluation of the area **(F)** and intensity **(G)** of P-gp immunolabeling. Arrowheads = endodermic epithelium; thin arrows = mesothelium; * = mesodermal blood vessels. Statistical differences were tested by Mann Whitney test. *p<0.05. Data are presented as mean ± SEM (n=6/group). Magnification bars represent 100 μm.

ABCG1 protein was detected in both the endodermic epithelium and in the endothelium of mesodermal vessels of the yolk sac, with lower levels of ABCG1 detected in the mesothelium. No ABCG1 was detected in the mesodermal layer connective tissue (Figures 4A and B). There were no differences in ABCG1 immunostaining area and intensity between PBA-infected and control pregnancies (Figures 4F and G).

**Fig. 4.**
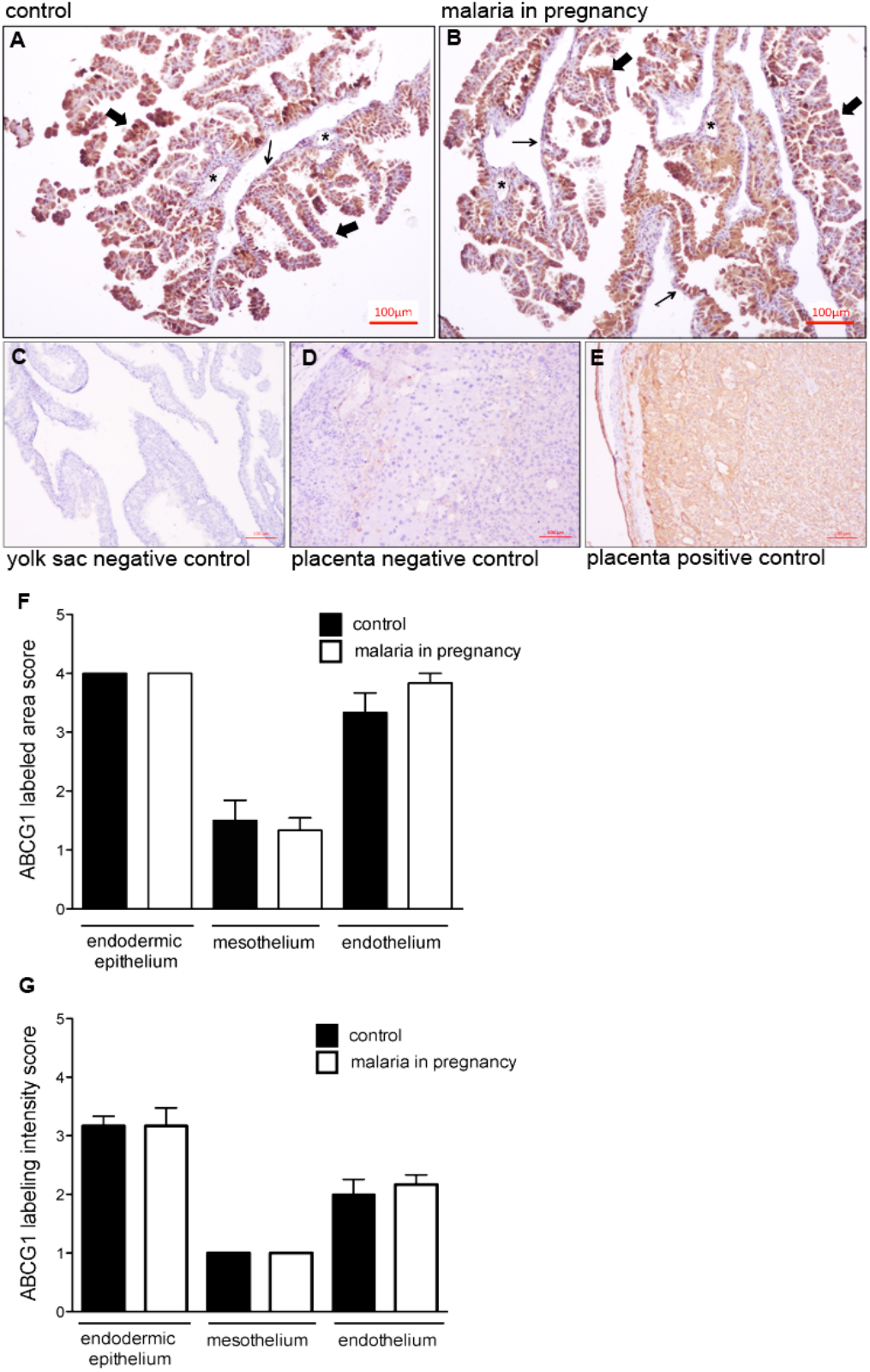
ABCG1 immunolocalization in the murine yolk sac. Representative immunohistochemistry images from murine yolk sac sections of control **(A)** and (**B)** *Plasmodium berghei ANKA*-infected dams at GD18.5. C-D: yolk sac **(C)** and placental **(D)** negative controls. **(E)** placental positive control showing ABCG1 immunoreactivity in labyrinthine and junctional zone cells. Semiquantitative evaluation of the area **(F)** and intensity **(G)** of ABCG1 immunolabeling. Arrowheads = endodermic epithelium; thin arrows = mesothelium; * = mesodermal blood vessels. Statistical differences were tested by Mann Whitney test. Data are presented as mean ± SEM (n=6/group). Magnification bars represent 100 μm.

A disconnect between mRNA and protein expression patterns for some ABC transporters have been previously reported in the placenta [16,34]. As such, we conducted immunohistochemical analysis of ABCA1 lipid and BCRP drug transporters in the mouse yolk sac. ABCA1 was primarily localized to the endodermic epithelium, with heterogeneous intensity. In addition, the cytoplasm of some epithelial cells was positive for ABCA1. The mesothelium and the endothelium of mesodermal vessels were also stained, but with less intensity compared to the endodermic epithelium (Figure 5A and B). No ABCA1 staining was observed in the connective tissue. Semi-quantitative analysis showed that both area and intensity of ABCA1 staining were increased in the endothelium of the mesodermal blood vessels (p <0.05) in MiP, with no changes in the endodermic epithelium or in the mesothelium (Figure 5F and G).

**Fig. 5.**
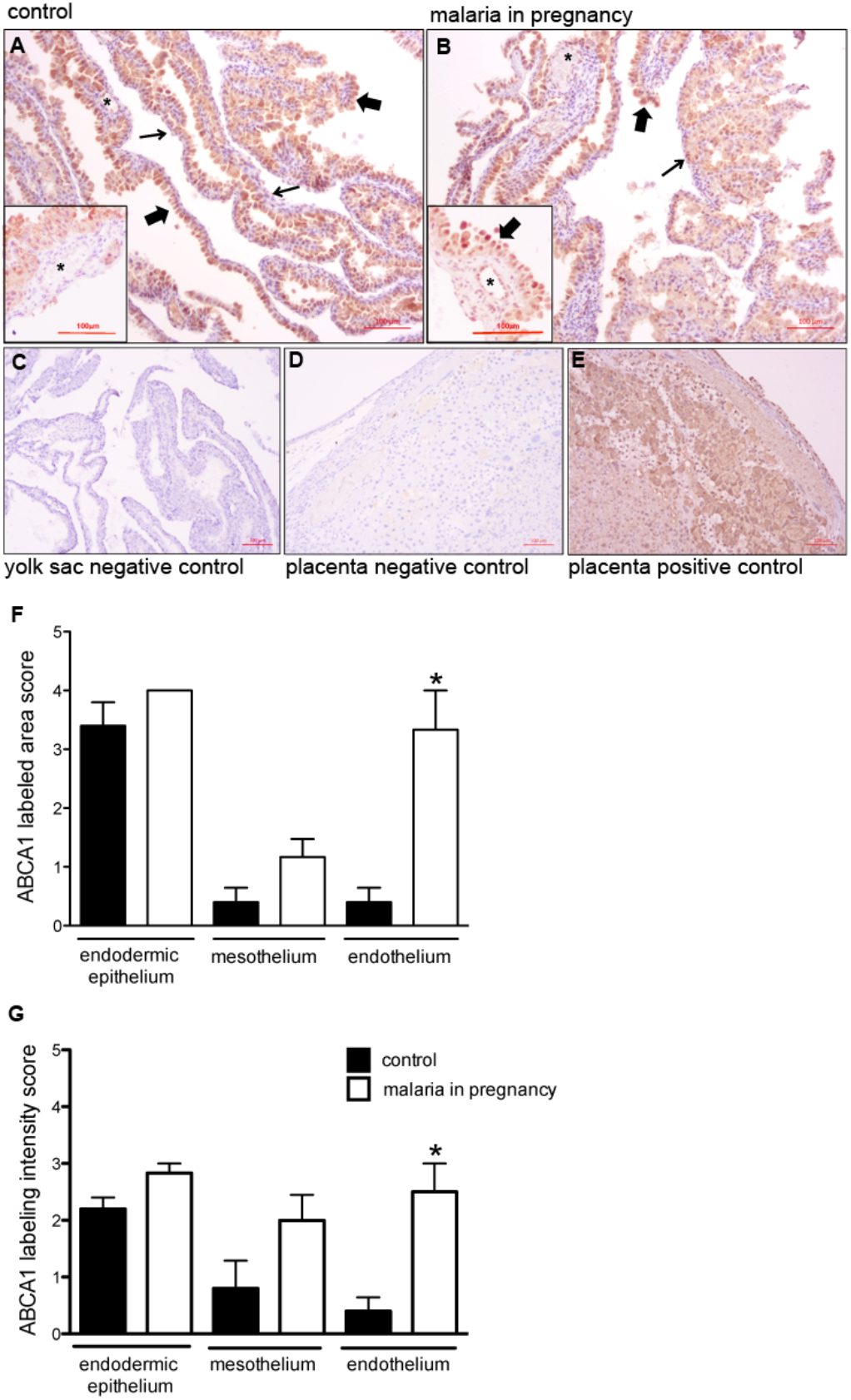
ABCA1 staining is increased in the endothelium of the mesodermal blood vessels in malaria-infected dams. Representative immunohistochemistry images from murine yolk sac sections of control **(A)** and **(B)** *Plasmodium berghei ANKA*-infected dams at GD18.5. Yolk sac **(C)** and placental **(D)** negative controls. **(E)** placental positive control showing ABCA1 immunoreactivity in labyrinthine and junctional zone cells. Semiquantitative evaluation of the area **(F)** and intensity **(G)** of ABCA1 immunolabeling. Arrowheads = endodermic epithelium; thin arrows = mesothelium; * = mesodermal blood vessels. Statistical differences were tested by Mann Whitney test. *p<0.05. Data are presented as mean ± SEM (n=6/group). Magnification bars represent 100 μm.

BCRP staining was localized to the endodermic epithelium and in the endothelium of mesodermal vessels, with lower levels in the mesothelium regions of the yolk sac. Semi-quantitative analysis revealed no effect of MiP on BCRP in any regions of the yolk sac (Figures 6A and B). As predicted, mouse placentae-positive controls exhibited intense ABCA1, ABCG1, BCRP and P-gp staining in labyrinthine and in junctional zone cells (Figure 3-6E), whereas negative controls showed minimal signal (Figure 3–6 C and D).

**Fig. 6.**
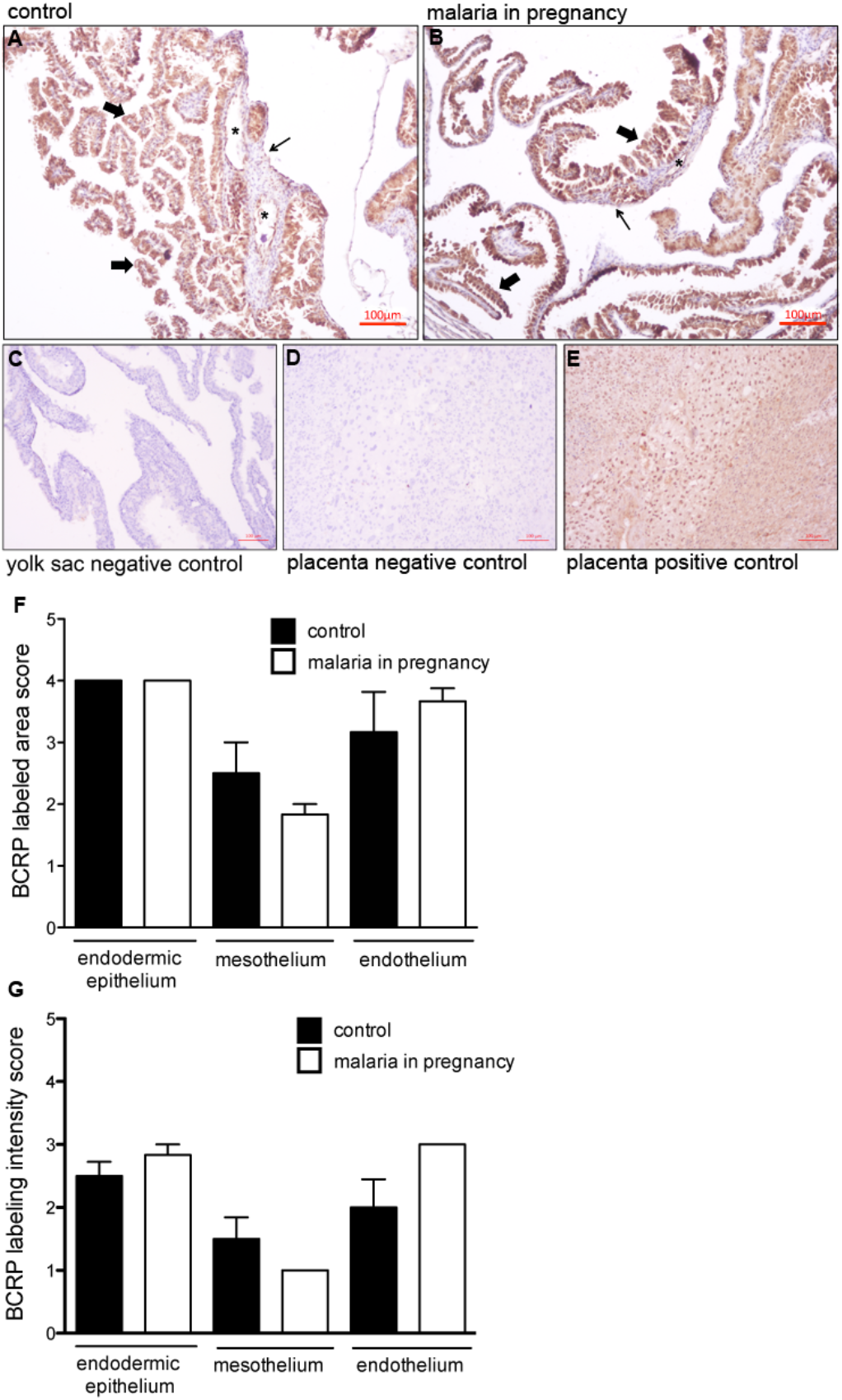
BCRP immunolocalization in the murine yolk sac. Representative immunohistochemistry images from murine yolk sac sections of control **(A)** and **(B)** *Plasmodium berghei ANKA*-infected dams at GD18.5. Yolk sac **(C)** and placental **(D)** negative controls. **(E)** placental positive control showing BCRP immunoreactivity in labyrinthine and junctional zone cells. Semiquantitative evaluation of the area **(F)** and intensity **(G)** of BCRP immunolabeling. Arrowheads = endodermic epithelium; thin arrows = mesothelium; * = mesodermal blood vessels. Statistical differences were tested by Mann Whitney test. Data are presented as mean ± SEM (n=6/group). Magnification bars represent 100 μm.

## Discussion

Our study provides new insights into how the yolk sac exerts embryo and fetal protection. We have also demonstrated how MiP impacts the expression of transporters that modulate yolk sac permeability to drugs, environmental toxins and nutrients. Using a mouse model of malaria induced-IUGR and PTL, we detected increased expression of P-gp/*Abcb1a* and *Mif*-chemokine in the yolk sac, and endothelial ABCA1 staining in blood vessels. In contrast, we demonstrated a decrease yolk sac *Abcg1* mRNA levels in MiP.

Previously, we have demonstrated that MiP impairs the placental expression of ABCA1/*Abca1*, P-gp/*Abcb1b* and BCRP/*Abcg2*, which were accompanied by increased placental levels of *Cxcl1* and *Ccl2* mRNA and upregulation of maternal *Il1-β, Il-6, Cxcl1* and *Ccl2* [9]. In the previous study, we provided evidence that MiP has the potential to increase fetal exposure to drugs and environmental toxins, via downregulation of major placental drug and nutrient ABC efflux transporters [9]. Moreover, other studies have demonstrated the sensitivity of placental P-gp and BCRP to infection. C57BL/6 mice exposed to sublethal (fetal) LPS in mid pregnancy, exhibited impaired placental P-gp activity [18], while polyinosinic:polycytidylic acid (PolyI:C, a viral mimic), had no effect on placental P-gp activity [35]. Human first trimester placentae exposed to lipopolysaccharide (LPS, a bacterial antigen), exhibited decreased *P-gp/ABCB1* and BCRP/*ABCG2* levels [36]. In contrast, human chorioamnionitis in 2^nd^ trimester, often resulting from polymicrobial infection [37], resulted in decreased P-gp levels but elevated BCRP expression [16]. Further, extravillous trophoblast (HTR8/SVneo)-like cells treated with LPS and single stranded RNA (ssRNA, another viral mimic), showed a profound downregulation of BCRP/*ABCG2* [14]. Together these studies demonstrate that the nature of the infective stimuli, determines the type of the drug efflux transporter response in different trophoblast lineages.

Notably, in the present study, we did not observe substantial up-regulation of pro-inflammatory factors in the yolk sac following MiP. Out of the five pro-inflammatory genes investigated, *Mif* chemokine was up-regulated, *Il-1 β* and *Ccl2* were unchanged and *Il-6* and *Cxcl1* were not detectible. In this context, the MIF chemokine is present at high levels in the amniotic fluid of women in PTL with infection and its expression is increased in chorioamniotic membranes during infection [38].

The pattern of the pro-inflammatory response observed in the yolk sac is in contrast with our previous report showing placental up-regulation of *Cxcl1* and *Ccl2*, induced by MiP [9]. Together, these results suggest that the yolk sac mounts a blunted pro-inflammatory response to systemic malarial infection when compared to the placenta. The placenta would appear to be efficient in buffering the transfer of malarial antigens and related pro-inflammatory cytokines/chemokines to the yolk sac and to the embryo/fetus. This may explain, at least in part, the different ABC transporter response pattern in the yolk sac, compared to the placenta.

In rodents, P-gp is encoded by two different gene isoforms, *Abcb1a* and *Abcb1b*. Yolk sac *Abcb1a* was the gene isoform up-regulated by MiP and was associated with increased P-gp immunostaining in both the uterine-facing membrane of endodermic cells and in the inner, amnion side-facing, mesothelial cells. While MiP did not alter the morphology of the yolk sac wall, the changes in P-gp and ABCA1, likely alter the barrier function of the yolk sac. However our morphometric analysis detected a higher number of constitutive outer endodermic cells (≈66%) in comparison to inner mesothelial cells (≈5%); in the control and MiP groups. This higher number of constitutive endodermic cells in the yolk sac, in concert with increased cell membrane P-gp, suggest that MiP leads to a greater net outflow of P-gp substrates from the fetal side, towards the uterine cavity (endoderm-mediated), rather than into the fetal side (mesothelium-mediated), likely favoring fetal protection.

Apart from drugs and environmental toxins, P-gp also transports pro-inflammatory compounds, Il-2, INF-γ, TNF-α and CCL2. As such, it can play an important role in the regulation of local inflammatory response [39,40]. Up-regulation of yolk sac Pgp in PBA-infected dams indicates this membrane has the potential to act together with other gestational tissues, such as the placenta, to participate in the immunomodulatory response to MiP. It is currently unknown whether this represents a compensatory mechanism to circumvent decreased placental P-gp and BCRP expression induced by MiP, or if this is an inherent response to MiP infection. This clearly requires further investigation.

Despite reduced yolk sac *Abcg1*, no changes in ABCG1 staining were observed. In contrast, increased ABCA1 levels were detected only in yolk sac endothelial cells of PBA-infected pregnancies. ABCA1 and ABCG1 are important cholesterol transporters, located on both the plasma and endosomal cell membranes [41]. In gestational tissues, apart from being localized in human syncytiotrophoblasts, ABCA1 and ABCG1 are also present in fetal endothelial cells of the placenta, acting in the exchange of cholesterol and phospholipids in the maternal interface [12,42]. Importantly, ABCA1 can also be localized in intracellular compartments such as the endoplasmic reticulum, functioning as regulator of intracellular signaling, cell differentiation, and hormone metabolism [43]. Changes in the expression of cholesterol transporters may adversely impact pregnancy outcome. Inhibition of ABCG1 and ABCA1 expression by siRNA transfection increased the sensitivity of human trophoblast cells to cytotoxic 25-hydroxycholesterol and 7-ketocholesterol oxysteroids, resulting from placental oxidative stress. On the other hand, induction of ABCA1 and ABCG1 in the trophoblast conferred placental protection against these agents [42], demonstrating an important role of these lipid efflux transporters protecting trophoblast cells against cytotoxic lipid derivates. Furthermore, disruption of ABCA1 and ABCG1 may be harmful to the establishment of pregnancy and placentation, since they regulate hormone synthesis and nutrient transport in the placenta [44]. Increased vascular ABCA1 may be the result of the yolk sac attempting to maintain normal cholesterol homeostasis in the fetal compartment, given that cholesterol is an essential element for the development and survival of the central nervous and other fetal organ systems [16,45].

In addition, we did not identify changes in gene expression of neutral amino acid transporters (*Snat1* and *Snat2*) or of *Snat4* and *Glut1* glucose transporters in the yolk sac in MiP. Such data suggest that gene expression of neutral amino acid and glucose transporters, at least in term pregnancies, is less impacted by malarial infection.

In conclusion, we have shown, for the first time, that yolk sac efflux transport potential may be disrupted by MiP. The presence of ABC transporters in the yolk sac wall and their selective alterations induced by MiP, suggest that this fetal membrane acts as an important protective gestational barrier under normal conditions as well as in malaria disease. Changes in the expression pattern of yolk sac ABCA1 and P-gp may alter the biodistribution of toxic substances, xenobiotics, nutrients and immunological factors within the fetal compartment and therefore participate in the pathogenesis of malaria induced-IUGR and PTL. In addition, based on our data, it is possible to hypothesize that MiP, along with other infective processes, has the potential to disrupt the human yolk sac protective barrier and thus impact early pregnancy outcome. This highlights the importance of investigating the human yolk sac response to infection.

## Acknowledgments

We thank Carlos Henrique da Silva for technical assistance and Drs. Cristina Fonseca Guatimosim and Patricia Massara Martinelli for assistance in imaging analysis and acquisition.

This study was supported by the Bill & Melinda Gates Foundation (MCTI/CNPq/MS/SCTIE/Decit/Bill and Melinda Gates 05/2013; OPP1107597), the Canadian Institutes for Health Research (SGM: Foundation-148368), Conselho Nacional de Desenvolvimento Científico e Tecnológico (CNPq; 304667/2016-1, 422441/2016-3, 303734/2012-4, 422410/2016-0); Coordenação de Aperfeiçoamento Pessoal de Nível Superior (CAPES, finance Code 001); Fundação de Amparo à Pesquisa do Estado do Rio de Janeiro (FAPERJ, CNE 2015/E26/203.190/2015); and PRPq-Universidade Federal de Minas Gerais (PRPq-UFMG, 26048).

